# 229E- and NL63-like coronaviruses in phyllostomid bats, Belize

**DOI:** 10.1101/2025.04.08.647854

**Authors:** B. R. Ansil, Guang-Sheng Lei, Aaron J. Santos, Michael Letko, Stephanie N Seifert, M. Brock Fenton, Nancy B. Simmons, Ryan F. Relich, Daniel J. Becker

## Abstract

We report novel and previously identified alphacoronavirus (α-CoV) diversity in three phyllostomid bat species (*Desmodus rotundus, Carollia sowelli*, and *Sturnira parvidens*) in Belize. Phylogenetic analysis suggests strong similarity between two Neotropical bat α-CoV lineages and human CoVs (229E and NL63) compared to prior bat α-CoVs.

## Introduction

Coronaviruses (CoVs), paramyxoviruses, and lyssaviruses are among the most epidemiologically relevant zoonotic viruses of bats (1). These viruses have caused outbreaks, epidemics, and even pandemics in humans and domestic animals, emphasizing the importance of understanding their diversity and prevalence in wild bats (1). However, most CoV and paramyxovirus studies have focused on bats from the Eastern Hemisphere, especially in the tropics (2,3). The Neotropics are under-sampled for both bat viruses, despite hosting the world’s most diverse bat communities (2) and showing evidence for zoonotic lineages (4). While substantial work has focused on rabies virus, studies are largely biased towards *Desmodus rotundus* (the common vampire bat) (5).

The Neotropics host the most diverse bat communities globally, with the family Phyllostomidae representing a unique radiation of >200 species spanning divergent ecologies (6). This diversity offers unique scenarios for viral diversification and specialization (7). Earlier studies have sampled Neotropical bats from Brazil, Costa Rica, and Mexico, identifying rabies virus, α-CoVs and β-CoVs, and rubulaviruses and morbilliviruses (*Paramyxoviridae*) (8). However, the diversity, geographic range, and host range of such viruses, and their clinical implications, remain mostly poorly understood across the Neotropics. Our study contributes to this effort by investigating the presence and diversity of these viral taxa (CoVs, paramyxoviruses, and lyssaviruses) and improving sampling efforts for this disproportionately underrepresented region.

### The study

In April 2019, we conducted preliminary viral surveillance among three phyllostomid bat species in Belize: *Desmodus rotundus* (n=27), *Carollia sowelli* (Sowell’s short-tailed bats; n=10), and *Sturnira parvidens* (little yellow-shouldered bats; n=11). *D. rotundus* is broadly distributed across the Neotropics, whereas the latter species are restricted to Central America (Figure 1A). We focused on these species based on prior reports of all three viral taxa in *D. rotundus* and from the genera *Carollia* and *Sturnira* (8). Over 12 nights (6–10 PM), we captured bats in the Lamanai Archeological Reserve, Orange Walk District. We used sterile miniature rayon swabs (1.98 mm, Puritan) and sterile cotton swabs (4.78 mm, Puritan) to collect saliva (OS) and rectal (RS) samples in 1 mL DNA/RNA Shield (Zymo Inc) (Table 1). We also collected paired swabs in viral transport medium (VTM; Rocky Mountain Biologicals) to attempt viral isolation. We opportunistically collected feces and urine in 1 mL DNA/RNA Shield. Samples were preserved in a cryoshipper prior to storage at −80°C. All bats were released after processing. Sampling was approved by the Institutional Animal Care and Use Committee of the American Museum of Natural History (AMNHIACUC-20190129) and Belize Forest Department (FD/WL/1/19(06), FD/WL/1/19(09)).

**Figure 1:**
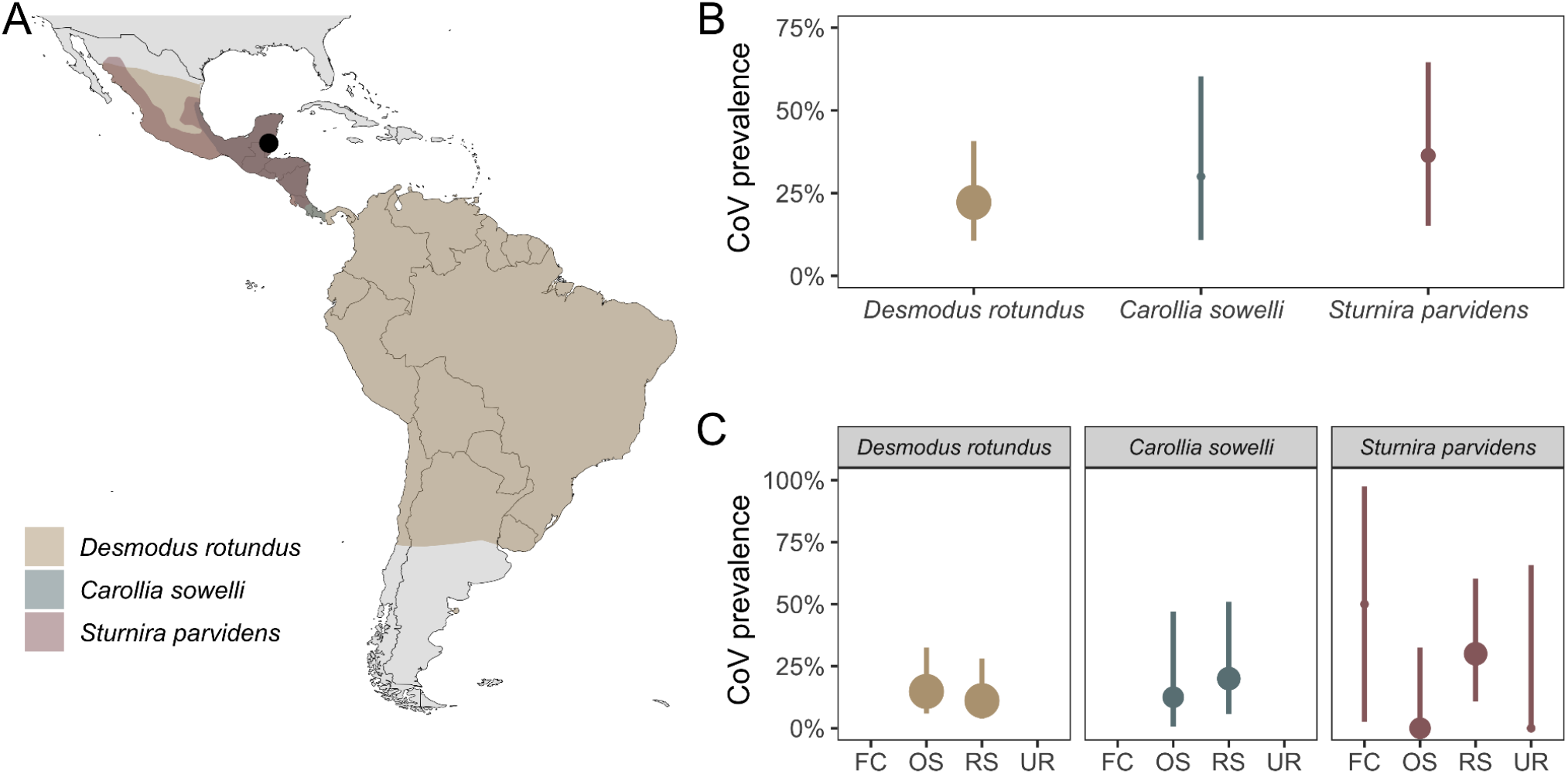
A) Map showing the sampling location in Belize (black point) with the distribution of *Desmodus rotundus, Sturnira parvidens*, and *Carollia sowelli*. B) Species-level CoV prevalence based on positivity in at least one sample type tested per individual. C) CoV prevalence by sample type: feces (FC), oral swabs (OS), rectal swabs (RS), and urine (UR). FC and UR were only collected from *S. parvidens*. Point estimates of CoV prevalence are scaled by sample size.

**Figure 2:**
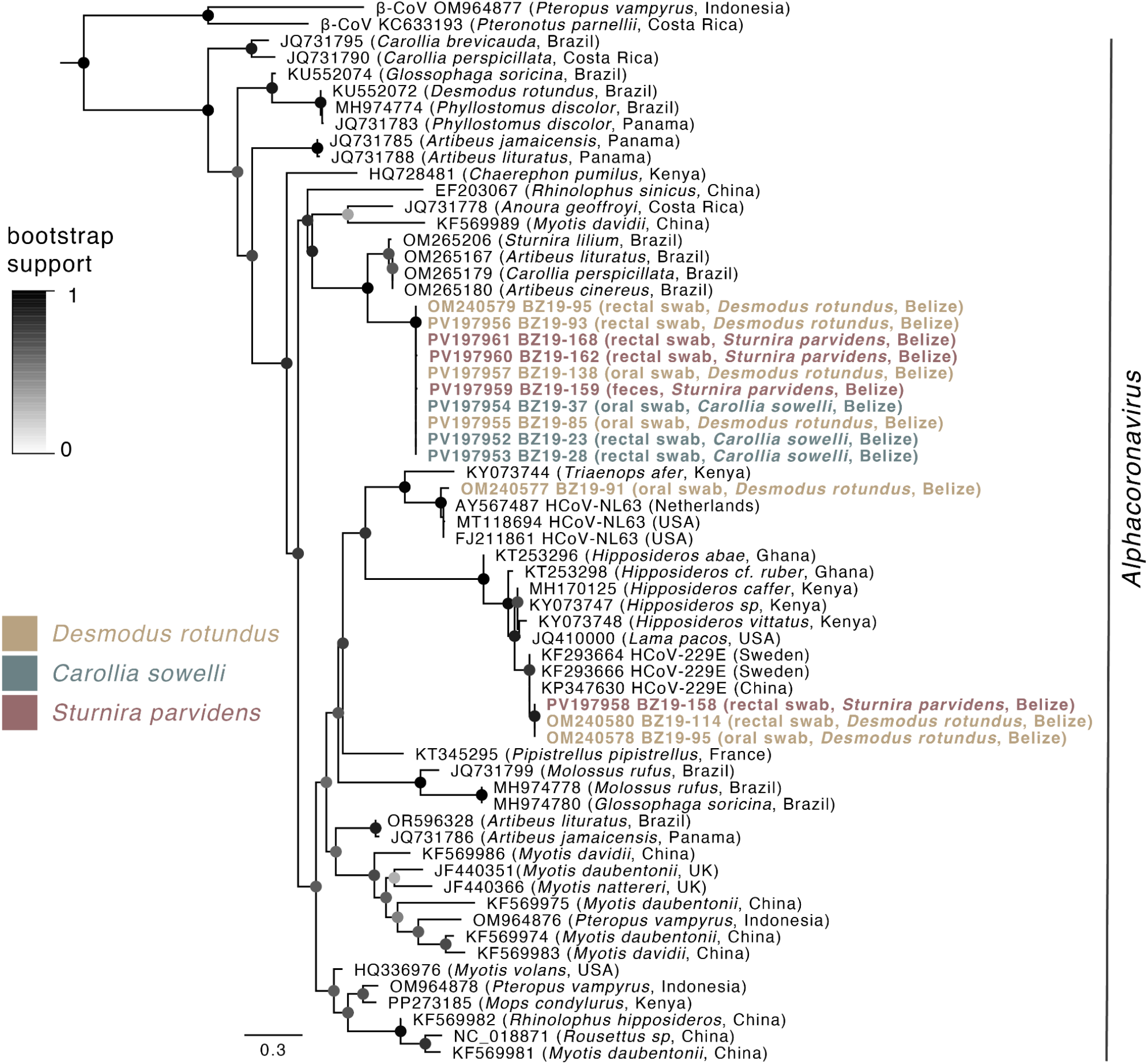
Maximum likelihood phylogeny displaying the evolutionary relationships among bat α-CoVs, HCoVs 229E and NL63, and the alpaca α-CoV using partial RdRp sequences. GTR+G nucleotide substitution model with 1,000 bootstrap replicates was used to infer tree topology and assess node support. The phylogeny was rooted with two bat β-CoV sequences. Sequences generated from Belize bats here are highlighted. Nodes are colored by bootstrap support (values <50% are not shown).

**Table 1.** Sample size distribution across the analyzed three bat species and four sample types.

|  | Saliva (OS) | Rectal swab (RS) | Feces (FC) | Urine (UR) |
| --- | --- | --- | --- | --- |
| *Desmodus rotundus* (n=27) | 27 | 27 | - | - |
| *Carollia sowelli* (n= 10) | 8 | 10 | - | - |
| *Sturnira parvidens* (n=11) | 8 | 10 | 2 | 2 |
| Total (n= 48) | 43 | 47 | 2 | 2 |

We extracted RNA from samples in DNA/RNA Shield using the *Quick*-RNA Viral Kit (Zymo Research). RNA was converted into cDNA using an iScript Synthesis Kit (BioRad), used as the template in three semi-nested PCRs for viral diagnostics. For CoVs, we targeted a 216-bp fragment of the RNA-dependent RNA polymerase (RdRp) gene (9). We targeted a 200-500 bp fragment of the L gene (10) and a 164-bp fragment of the N gene (11) for paramyxoviruses and lyssaviruses, respectively. All PCRs included negative controls. Amplicons were Sanger sequenced and aligned using Geneious, followed by analysis using NCBI BLAST and deposition in GenBank (PV197952−61). A subset (n=4) of CoV sequences were published previously (OM240577−80) (12).

We detected CoVs in all three species with varying prevalence (Figure 1B): *D. rotundus* (22.22%, 95% confidence interval [CI]:10.6–40.75%), *C. sowelli* (30%, 95% CI: 10.77– 60.32%) and *S. parvidens* (36.36%, 95% CI: 15.16–64.61%). We also observed variable prevalence between sample types within species (Figure 1C). In *D. rotundus*, OS had higher prevalence (14.81%, 95% CI: 5.91–32.47%) compared to RS (11.11%, 95% CI: 3.85–28.05%). In *C. sowelli*, RS had higher prevalence (20%, 95% CI: 5.66–50.98%) compared to OS (12.5%, 95% CI: 0.64–47.08%). For *S. parvidens*, we had urine and feces as additional samples, and feces showed higher prevalence (50%, 95% CI: 2.56–97.43%) than RS (30%, 95% CI: 10.77– 60.32%); no OS (n=8) or urine (n=2) samples were positive. We fit a generalized linear model with binary response and small sample size bias correction in R with the *brglm2* package to assess the effects of species, sample type (OS or RS) and their interaction on the probability of infection. However, a likelihood ratio test revealed no significant effect of species (*χ*^*2*^*=*0.20, *P*=0.90), sample type (*χ*^*2*^*=*2.36, *P*=0.49), or their interaction (*χ*^*2*^*=*1.96, *P*= 0.37) on CoV infection status. No samples were positive for paramyxoviruses or lyssaviruses.

We aligned our CoV RdRp sequences, along with other bat and human CoVs, using MUSCLE in Geneious, and constructed a maximum-likelihood phylogeny using PhyML in NGPhylogeny.fr. Our phylogeny revealed three distinct α-CoV lineages in Belize phyllostomids. The first lineage is novel and consists of ten monophyletic sequences from all three bat species, forming a sister lineage to α-CoV sequences reported from other phyllostomids in Brazil (e.g., OM265179–80). This distribution suggests strong local-scale circulation among host species while maintaining evolutionary similarity with congeners in the Neotropics. Notably, the other two lineages show close phylogenetic similarity to NL63 and 229E, two human coronaviruses (HCoV) associated with respiratory tract infections (13). Specifically, the second lineage, previously identified and consisting of a single sequence from *D. rotundus* (OM240577) (12), is phylogenetically similar to multiple lineages of NL63 (e..g., MT118694), with a NL63-like bat CoV from Kenya forming a basal clade. The third lineage included sequences previously identified from two *D. rotundus* (i.e., OM240580, OM240578) (12) and one newly identified from *S. parvidens* (PV197958), both exhibiting close phylogenetic similarity to 229E (e.g., KP347630); all other 229E-like bat CoVs formed a basal clade to this group. The alpaca α-CoV (JQ410000), included due to its evolutionary relationship with 229E, clustered with bat CoV sequences from the Afrotropics but not these Neotropical bat 229E-like CoVs.

For PCR-positive samples, we attempted viral isolation by inoculating eluates from paired VTM samples onto monolayers of Vero E6 cells (ATCC CRL-1586) and primary kidney, brain, lung, liver, and spleen primary cell cultures derived from *Carollia perspicillata* (14). We did not detect viral infection–induced cytopathic effects nor CoV RdRp amplicons in any of the cultured samples during a 20-day period. Viral culture was performed under BSL-3 conditions at Indiana University (Institutional Biosafety Committee protocol IN-1348).

## Conclusions

Our study provides significant insights into the diversity and circulation of CoVs in the Neotropics, identifying three distinct α-CoV lineages in bats from Belize. One lineage was novel and shared between *D. rotundus, C. sowelli*, and *S. parvidens*, suggesting strong local-scale circulation and multiple host-switching events. The other lineages were closely related to HcoVs while corroborating the known evolutionary ancestry of 229E to Afrotropical insectivorous bats (15). However, the similarity of these bat α-CoV sequences to HCoVs compared to Afrotropical bat sequences poses several questions, including the directionality of spillover (bats to humans or humans to bats) and risk to both bats and humans. Addressing these questions will require surveillance targeting viral genomes from a broad range of host taxa across the wider Neotropics.

## Acknowledgements

For assistance with bat sampling logistics and research permits, we thank Mark Howells, Neil Duncan, and the staff of the Lamanai Field Research Center. This work was supported by the National Geographic Society (NGS-55503R-19), Edward Mallinckrodt, Jr. Foundation, Human Frontier Science Program (RGP002/2023, LT0017/2024-L), Indiana University, and the American Museum of Natural History (Taxonomic Mammalogy Fund).

## Data availability

All RdRp sequence data are available on GenBank under accessions OM240577−80 and PV197952−61.

## Conflicts of interest

The authors declare no conflicts of interest.

## Notes

### Competing Interest Statement

The authors have declared no competing interest.

